# From culture to clarity in four hours: accelerating clinical management of bloodstream infections using metagenomics

**DOI:** 10.64898/2026.07.20.739511

**Authors:** Jawad Ali, Anurag Basavaraj Bellankimath, Silje Therese Opgård, Emil Varman Manivannan, Gunnar Skov Simonsen, Rafi Ahmad

## Abstract

**Background:** Metagenomic next-generation sequencing (mNGS) has the potential to transform clinical diagnostics for bloodstream infections (BSIs). However, its clinical utility is currently limited by several challenges, including the extraction of DNA from blood cultures. Our aim was to develop, evaluate, and optimize an in-house method for host depletion and bacterial DNA extraction from positive blood cultures to enable rapid mNGS-based pathogen detection and antimicrobial resistance profiling, informing clinical management of BSIs.

**Methods:** 151 clinical blood cultures (115 positive and 36 negative) were processed for DNA extraction using an in-house-developed SEPSINN method for host depletion and bacterial DNA extraction. mNGS on the MinION was performed, and the results for pathogen identification and antimicrobial susceptibility predictions were compared with the routine clinical workflow. SEPSINN was also evaluated against a commercial DNA extraction method to assess its effectiveness in depleting host DNA and recovering bacterial DNA.

**Results:** The SEPSINN method achieved up to 1000-fold depletion of host DNA and outperformed the commercial DNA extraction method. At the sample level, mNGS achieved 100% accuracy, specificity, and sensitivity, identifying at least one pathogen in all 115 positive blood cultures. At the pathogen level, mNGS showed 98% accuracy (120/123), specificity, and sensitivity. For antimicrobial susceptibility predictions, mNGS achieved an accuracy of 95% (1382/1451), a sensitivity of 88% (203/230), and a specificity of 97% (1179/1221). Moreover, the method also identified fungi, indicating a wider taxonomic range. mNGS resulted in an approximately 4-hour turnaround time for pathogen identification and resistance profiling.

**Conclusions:** The method can provide information on BSI clinical management within approximately 24 hours of receiving the sample, including the time required for culture positivity. This represents an important advancement in the clinical management of BSIs, with the potential to save lives and promote antibiotic stewardship.

## Background

Bloodstream infections (BSIs) and sepsis are a global health problem, responsible for millions of deaths each year [1,2]. In BSIs, bacteria are the predominant causative agents, accounting for around 90% of infections, while fungi, viruses, and other microorganisms account for roughly 10% of cases [3]. Early diagnosis and appropriate antibiotic administration are crucial for reducing mortality. Current diagnostics rely on blood culture, followed by matrix-assisted laser desorption/ionization time-of-flight mass spectrometry (MALDI-TOF MS) for pathogen identification and automated or manual antibiotic susceptibility testing [4]. However, these methods require 48 to 72 hours, which is crucial for septic patients, as every hour’s delay in antimicrobial treatment decreases survival by 7.6% [5,6]. Antimicrobial resistance (AMR) presents a unique challenge by complicating the selection of appropriate empirical therapy, underscoring the importance of rapid diagnostics.

Molecular diagnostic methods, such as multiplex PCR, offer a promising alternative because they are fast and can provide results within 1–4 hours after a positive blood culture [7]. However, multiplex PCR panels are limited to identifying only certain species and resistance determinants [8,9]. Metagenomic next-generation sequencing (mNGS) can overcome the limitations of multiplex PCR panels by identifying pathogens and resistance markers in a sample without relying on predefined targets [10,11]. Previous studies have demonstrated the potential of mNGS to detect pathogens and antibiotic resistance genes (ARGs) in blood culture [12–16]. Govender *et al.* reported 97% sensitivity and 94% specificity for pathogen identification using mNGS compared with routine matrix-assisted laser desorption ionization time-of-flight (MALDI-TOF) diagnostics [12]. Harris et al. have reported 94% accuracy in species identification using nanopore sequencing compared with routine diagnostics [16].

While mNGS can rapidly and accurately identify pathogens and detect ARGs in blood cultures, it faces some challenges. One of the main challenges is extracting pure microbial DNA, as blood cultures contain a high abundance of host cells and contaminants, including sodium polyanethol sulfonate (SPS), hemoglobin, and heme [17–19]. Therefore, there is an urgent need to develop a method that depletes host cells, removes inhibitors, and extracts high-quality bacterial DNA for mNGS-based workflows. Previous studies have employed selective host cell lysis with detergents such as saponin, followed by digestion with endonucleases to remove host DNA and extract bacterial DNA [20–23]. However, these studies were focused on clinical samples from respiratory and urinary tract infections and spiked blood cultures. In our previous study, we developed a saponin- and nuclease-based host DNA-depleting method for blood cultures [24]. However, the method was developed and primarily validated using spiked blood cultures.

In this study, we aimed to modify, evaluate, and optimize our in-house-developed host depletion method for use in clinical blood cultures [24]. It was followed by real-time nanopore sequencing and data analysis to identify pathogens and antimicrobial susceptibility profiling. The results from mNGS were used to assess concordance with routine MALDI-based species identification and disk diffusion assay for antimicrobial susceptibility testing (AST). The in-house-developed method achieved up to 1000-fold depletion of host DNA compared with non-depleted control samples. The mNGS-based method outperformed the *initial MALDI* for pathogen identification and performed on par with the *final MALDI*, achieving 98% accuracy, specificity, and sensitivity. Similarly, for AST predictions, mNGS achieved 95% accuracy, 88% sensitivity, and 97% specificity compared with routine disk diffusion testing.

## Methods

### Sample collection and routine microbiological testing

This study involved the collection of 151 blood cultures at the University Hospital of North Norway (UNN). It included 115 consecutive positive blood cultures from adults and 36 randomly chosen negative blood cultures. The routine diagnostic laboratory used aerobic and anaerobic BACT/ALERT^®^ Culture Media Bottles (bioMérieux, Marcy-l’Étoile, France) for culturing, followed by incubation in the BACT/ALERT^®^ VIRTUO^®^ system. The microbiology laboratory at UNN performed two different MALDI analyses. The first MALDI, referred to as "*initial MALDI*", is performed directly from positive blood cultures according to the manufacturer’s instructions. A second MALDI, hereafter referred to as "*definite MALDI*", is subsequently performed on purified bacterial colonies obtained after subculturing. Antibiotic susceptibility testing (AST) was performed using the EUCAST disk diffusion method, and the results were reported as resistant (R), susceptible to standard exposure (S), or susceptible to increased exposure (I) according to the EUCAST breakpoint protocol v16.0 [25]. These routine results for pathogen identification and AST were used for benchmarking mNGS-based predictions.

### Host depletion and microbial DNA extraction

Host depletion and microbial DNA extraction were performed using an in-house-developed method called "SEPSINN". 1-2 mL of each blood culture sample was processed for host depletion as reported, with some modifications [24]. The blood culture samples were centrifuged at 13000g for 5 min, and the supernatant was discarded while the pellets were processed further. 600 µL of 8% saponin and 600 µL of HL-SAN buffer were added to obtain a final saponin concentration of 4%, followed by the addition of 250 U of HL-SAN endonuclease (25 U/µL, ArcticZymes Technologies, Tromsø, Norway). The sample was gently mixed and incubated at 37 °C for 15 min with shaking at 900 revolutions per minute. After incubation, the mixture was centrifuged to remove debris, and the pellet was washed with phosphate-buffered saline. Lysis of bacterial cells was performed using a bead-beating step with lysis buffer, followed by magnetic bead-based extraction with the NAxtra™ 2.0 Aquaculture NA extraction kit (Lybe Scientific AS, Trondheim, Norway).

### qPCR-based assays

To assess the host depletion efficiency of the SEPSINN method, DNA was extracted from 10 blood cultures (*E. coli* and *P. aeruginosa*) with and without host depletion. qPCR was performed on these samples to estimate host and bacterial DNA, targeting the human β-actin gene and the respective species-specific bacterial genes (Additional File 1: Table S1). The qPCR reactions were performed using a QuantStudio™ 5 instrument (ThermoFisher Scientific) under the same conditions as previously reported [24]. Each PCR reaction included 3 μL of 5X HOT FIREPol® EvaGreen® qPCR Supermix (Catalog no. 08-36-00001, Solid BioDyne, Estonia), 0.2 μM of forward and reverse primers, 10.4 μL of nuclease-free water, and 1 μL of template DNA. The qPCR amplification conditions were as follows: an initial denaturation at 95 °C for 12 min, followed by 40 cycles of 95 °C for 25 s, 60 °C for 45 s, and 72 °C for 60 s, with a final dissociation stage. Host depletion and bacterial DNA enrichment were assessed using ΔCt values, normalized to the non-depleted biological controls [23,24].

In addition, 12 blood culture samples from the most common BSI-causing pathogens, including *E. coli*, *S. aureus*, *K. pneumoniae*, *P. aeruginosa*, *S. pneumoniae*, *E. faecalis*, *S. epidermidis*, *S. marcescens*, and *B. fragilis,* were also processed with the QIAamp BiOstic Bacteremia DNA Kit (Qiagen, Germany). qPCR using species-specific primers (Additional File 1: Table S1) was performed to compare host depletion and bacterial DNA recovery between SEPSINN and BiOstic. Quantification of all extracted DNA samples was performed using the Qubit 4 fluorometer with the Qubit 1X dsDNA HS Assay Kit (Catalog no. Q33231, ThermoFisher Scientific). The purity of the extracted DNA was assessed using a Nanodrop ND-1000 spectrophotometer (ThermoFisher Scientific), which measured absorbance ratios at 260/280 nm and 260/230 nm.

### Sequencing and bioinformatics analysis

Library preparation for nanopore sequencing was performed using the Rapid Barcoding Kit 96 V14 (SQK-RBK114.96), with sequencing carried out on the R10.4.1 MinION flow cell (FLO-MIN114). The samples were multiplexed, with 20-30 loaded onto the flow cell per run. The sequencing data were basecalled in real time using the ONT MinKNOW GUI (Version 6.0.11), and high-accuracy basecalling was performed with the Dorado basecaller (Version 0.6.4). Unclassified reads were recovered using MysteryMaster [26]. The bioinformatic analysis was performed to identify pathogens and predict antibiotic susceptibility using an in-house-developed pipeline, as reported [23]. Briefly, the basecalled sequencing data were BLAST searched against a custom reference database of the most common sepsis-causing pathogens. Reads that did not align with any prokaryotic genome were presumed to be host reads and were discarded. If pathogen genome coverage exceeded 2%, reference-based assemblies were created using minimap2 [27] (version 2.22) with the “map-ont” option. Samtools [28] (version 1.13) was used to filter the alignments (options -b -F 4) and generate a consensus sequence. ARGs were identified using abricate [29] (version 1.0.1) against the NCBI [30] and the Comprehensive Antibiotic Resistance Database (CARD) [31] databases with default settings. Additionally, for false-negative predictions, K-MARVEL [32] was used to detect ARGs and identify relevant mutations. Multilocus sequence typing (MLST) analysis was performed on pathogens uniquely identified by mNGS using the OmicsBox (version 3.5.4) implementation of PubMLST [33].

### Statistical analysis

Accuracy, sensitivity, and specificity for mNGS predictions compared to the routine workflow were calculated at the sample, pathogen, and antibiotic levels.

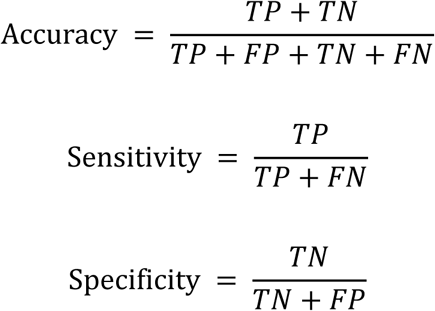

At the sample level, true positives (TP) are those that are flagged positive by both *definite MALDI* and mNGS. Similarly, true negatives (TN) were samples that were flagged negative by both. False positives (FP) were samples that were negative by *definite MALDI*, but mNGS showed as positive. False negatives (FN) are samples detected as positive by *definite MALDI* but negative by mNGS. A similar classification was performed at the pathogen level, with detection or non-detection considered for each pathogen rather than each sample. Similarly, for the AST predictions, any pathogen-antibiotic combination identified as susceptible (S) by both mNGS and the routine method was classified as TN, whereas those identified as resistant (R) by both methods were classified as TP. FPs were antibiotic-pathogen combinations for which mNGS showed resistance, but the routine method showed susceptibility, and FNs were those for which the routine method showed resistance while mNGS flagged them as susceptible.

To assess SEPSINN performance relative to the BiOstic kit, a Kruskal-Wallis H-test [34] was performed on the qPCR Ct values. A post hoc analysis – Dunńs test [35] was also conducted on this data set to compare the extraction methods. The analysis was performed using GraphPad Prism (Version 10.6.1), with a *p < 0.05* considered statistically significant.

A detailed overview of the experimental design is illustrated in Additional File 1: Fig S1.

## Results

### Sample diversity and routine microbiological results

Microbiological analysis identified 44 unique species, including 42 bacterial species (eight of which were anaerobic) and two *Candida spp*. (Additional File 1: Fig S2a). The time-to-positivity (TTP) for blood cultures ranged from 7 to 132 hours, with a mean of approximately 22 hours (Additional File 2: Table S2). The shortest mean TTP was around 17 hours for aerobic blood cultures. *Streptococcus dysgalactiae* and *Streptococcus agalactiae* had the shortest TTP, around 10 hours, followed by *Escherichia coli, Enterococcus faecalis,* and *Streptococcus pneumoniae,* with an average TTP of about 12 hours (Additional File 2: Table S2). The fungal species exhibited a slightly longer mean TTP of around 28 hours. The longest mean TTP (≈66 hours) was observed in the anaerobic blood culture, including for species *Cutibacterium acnes* (≈125 hours), *Eubacterium limosum* (81 hours), *Propionimicrobium lymphophilum* (74 hours), *Eggerthella lenta* (≈66 hours), *Fusobacterium nucleatum* (≈59 hours), and *Parvimonas micra* (≈43 hours).

In culture-positive samples, 95% (108/115) were monomicrobial, while 6% (7/115) were polymicrobial. Of the polymicrobial samples, six contained two species, and one contained three species (Additional File 1: Fig S2b). A total of 123 pathogens were identified through routine microbiological testing. *E. coli* was the most frequently detected pathogen, representing 23% (28/123) of all identified species. The second most prevalent pathogen was *S. aureus*, at 12% (15/123). Coagulase-negative staphylococci (CoNS) were also frequently detected, accounting for 21% (26/123) of all detected species. Three samples contained fungi, including two with *Candida albicans* and one with *Candida glabrata* (Additional File 1: Fig S2a).

### SEPSINN achieved effective host depletion and recovery of microbial DNA

The qPCR results demonstrated effective host depletion using the SEPSINN method compared with non-depleted samples. Overall, the method achieved 10^2^–10^3^-fold host depletion (Additional File 1: Table S3). *E. coli* samples 3 and 9 showed less depletion than the others, with ΔCt values of 5.82 and 0.6, respectively. The highest depletion was observed in sample 14 (*E. coli*), having a ΔCt of 9.7. Overall, most samples showed a 10^2^-fold depletion. Similarly, for the majority of samples, bacterial DNA recovery was higher in the depleted samples compared to their non-depleted counterparts. Only *E. coli* samples 5 and 9 showed a minor loss of bacterial DNA in the depleted samples (ΔCt = 2.1 and 1.6, respectively) compared to the non-depleted samples (Additional File 1: Table S3).

Moreover, compared with the BiOstic kit, SEPSINN efficiently depleted host DNA and recovered bacterial DNA. SEPSINN consistently outperformed BiOstic in host DNA depletion, with mean Ct values >40 for most samples (Additional File 1: Fig S3 and Table S4). The BiOstic kit showed host Ct values between 30 and 38, indicating higher host DNA levels than SEPSINN. Statistical analysis showed significant differences in host DNA depletion between SEPSINN and BiOstic, with a p-value of 0.002 and an H-score of 22.6 (Additional File 1: Fig S3d). For bacterial DNA, SEPSINN showed comparable or better recovery than BiOstic (Additional File 1: Fig S3). Similarly, for DNA obtained after extraction, SEPSINN yielded lower amounts than BiOstic in some samples (Additional File 1: Table S4). The lower yield resulted from SEPSINN’s efficient depletion of host DNA. However, the DNA yield was more than sufficient for downstream sequencing and qPCR analysis.

### mNGS-based pathogen identification outperformed the *initial MALDI* and was on par with *definite MALDI*

Nanopore sequencing data were acquired for each sample and correlated with the routine MALDI for pathogen detection (Additional File 2: Table S5). The mNGS method closely matched the *definite MALDI* for pathogen identification. However, it outperformed the *initial MALDI* for pathogen ID. The *initial MALDI* failed to identify or misidentified pathogens in 11 samples that mNGS correctly identified (Table 1). In samples 21, 26, 77, and 99, *initial MALDI* failed to identify the pathogens *S. capitis*, *S. aureus*, *E. coli*, and *E. faecalis*, respectively, whereas mNGS detected them. In samples 69 and 81, the *initial MALDI* incorrectly identified *Campylobacter jejuni* as *Starkeya novella* and *Bifidobacterium breve* as *Fannyhessea vaginae*, respectively. In sample 71, it failed to distinguish between *Neisseria meningitidis* and *Eikenella corrodens* (the correct pathogen). Moreover, the *initial MALDI* failed to detect pathogens in samples 80 (*Propionimicrobium lymphophilum*), 82 (*Parvimonas micra*), 83 (*Staphylococcus epidermidis*), and 96 (*Niallia circulans*). The *definite MALDI* was used as a reference to compare the performance of the *initial MALDI* and mNGS.

**Table 1.** Comparison of the species identification in samples in which the *initial MALDI* failed or incorrectly identified species compared to the reference, *definite MALDI,* and mNGS. Also, additional species identified by mNGS compared to the initial and definitive MALDI analyses are shown. *additional species detected by mNGS that were confirmed with qPCR and culturing on agar plates.

| Sample ID | Initial MALDI | Definite MALDI (Reference) | mNGS ID |
| --- | --- | --- | --- |
| 21 | *Staphylococcus haemolyticus* | *Staphylococcus haemolyticus* | *Staphylococcus haemolyticus* |
|  |  | *Staphylococcus capitis* | *Staphylococcus capitis* |
| 26 | *Staphylococcus epidermidis* | *Staphylococcus epidermidis* | *Staphylococcus epidermidis* |
|  |  | *Staphylococcus aureus* | *Staphylococcus aureus* |
| 69 | *Starkeya novella* | *Campylobacter jejuni* | *Campylobacter jejuni* |
| 71 | *Neisseria meningitidis/Eikenella corrodens* | *Eikenella corrodens* | *Eikenella corrodens* |
| 77 | *Streptococcus gallolyticus* | *Streptococcus gallolyticus* | *Streptococcus gallolyticus* |
|  |  | *Escherichia coli* | *Escherichia coli* |
| 80 | NO ID | *Propionimicrobium lymphophilum* | *Propionimicrobium lymphophilum* |
| 81 | *Fannyhessea vaginae* | *Bifidobacterium breve* | *Bifidobacterium breve* |
| 82 | NO ID | *Parvimonas micra* | *Parvimonas micra* |
| 83 | NO ID | *Staphylococcus epidermidis* | *Staphylococcus epidermidis* |
| 96 | NO ID | *Niallia circulans* | *Niallia circulans* |
| 99 | *Enterobacter hormaechei* | *Enterobacter hormaechei* | *Enterobacter hormaechei* |
|  |  | *Enterococcus faecalis* | *Enterococcus faecalis* |
| Additional species identified using mNGS |  |  |  |
| 1 | *Escherichia coli* | *Escherichia coli* | *Escherichia coli, Staphylococcus aureus\** |
| 2 | *Escherichia coli* | *Escherichia coli* | *Escherichia coli, Staphylococcus aureus\** |
| 5 | *Escherichia coli* | *Escherichia coli* | *Escherichia coli, Klebsiella pneumoniae* |
| 18 | *Escherichia coli* | *Escherichia coli* | *Escherichia coli, Staphylococcus aureus\** |
| 39 | *Escherichia coli* | *Escherichia coli* | *Escherichia coli, Staphylococcus aureus\** |

Compared to *definite MALDI*, mNGS achieved 100% accuracy, sensitivity, and specificity for sample-level identification (Fig 1a). mNGS identified at least one microbial species in all 115 positive samples. At the pathogen level, mNGS showed 98% accuracy, sensitivity, and specificity (Fig 1b and Additional File 1: Table S6). Out of the 123 pathogens identified by *definite MALDI*, mNGS correctly identified 120 species. The method failed to detect three pathogens: *Pseudomonas aeruginosa* in sample 45*, C. glabrata* in sample 52, and *Achromobacter xylosoxidans* in sample 108 (Fig 2). All three of these missed pathogens were from polymicrobial samples in which other pathogens were correctly identified. The *definite MALDI* could not distinguish between *Streptococcus oralis* and *Streptococcus mitis* in sample 56, but mNGS identified it as *S. oralis*. However, mNGS also detected *S. pneumoniae* in the same sample.

**Fig 1:**
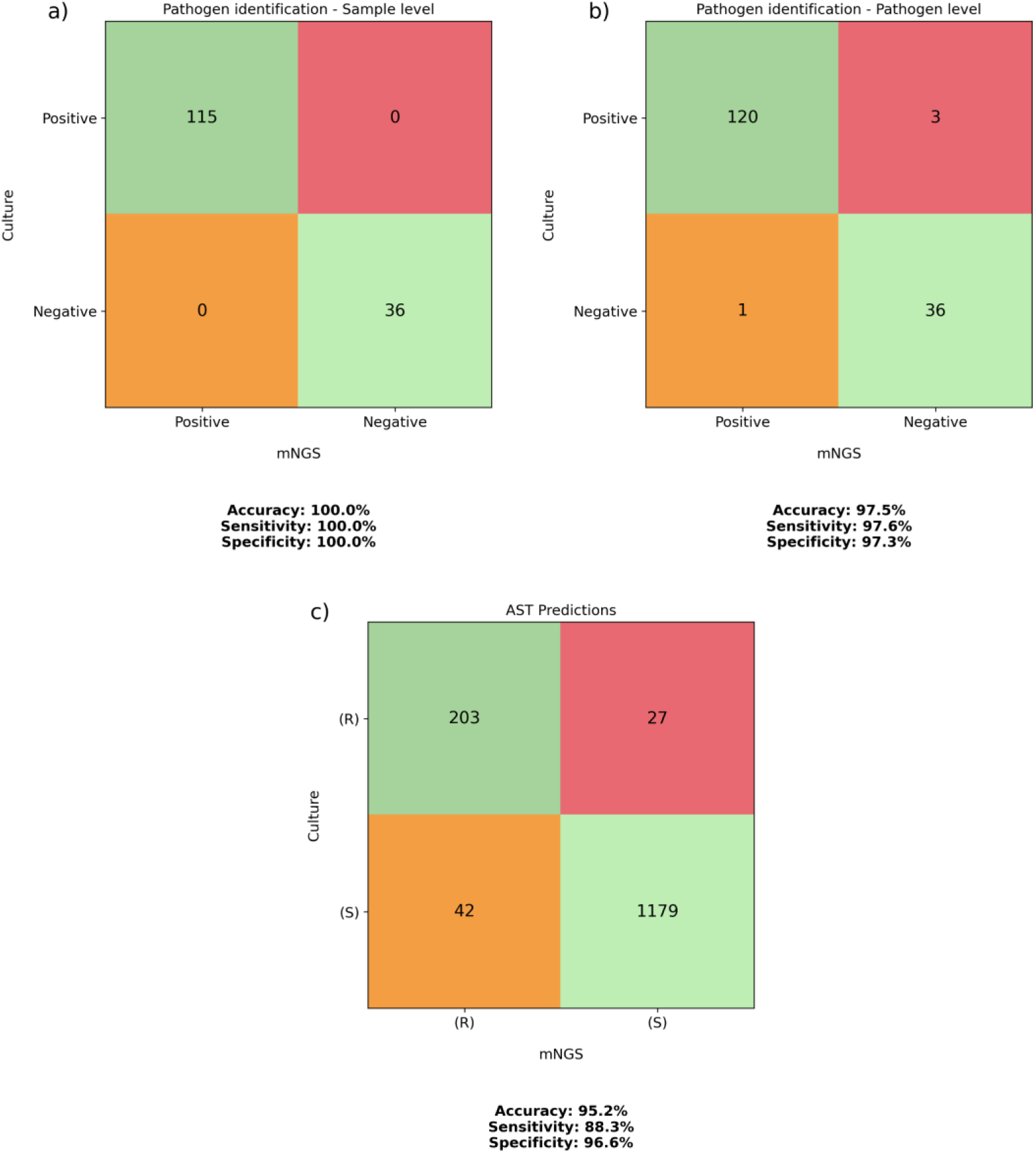
Confusion matrix showing the performance of the SEPSINN mNGS method for pathogen identification and antimicrobial susceptibility predictions. The *definite MALDI* was used as a reference for species identification, and disk diffusion for AST predictions. Subfigures (a) & (b) represent the performance of mNGS identification at the sample and pathogen level, while subfigure (c) shows the antimicrobial susceptibility predictions at the antibiotic level.

**Fig 2:**
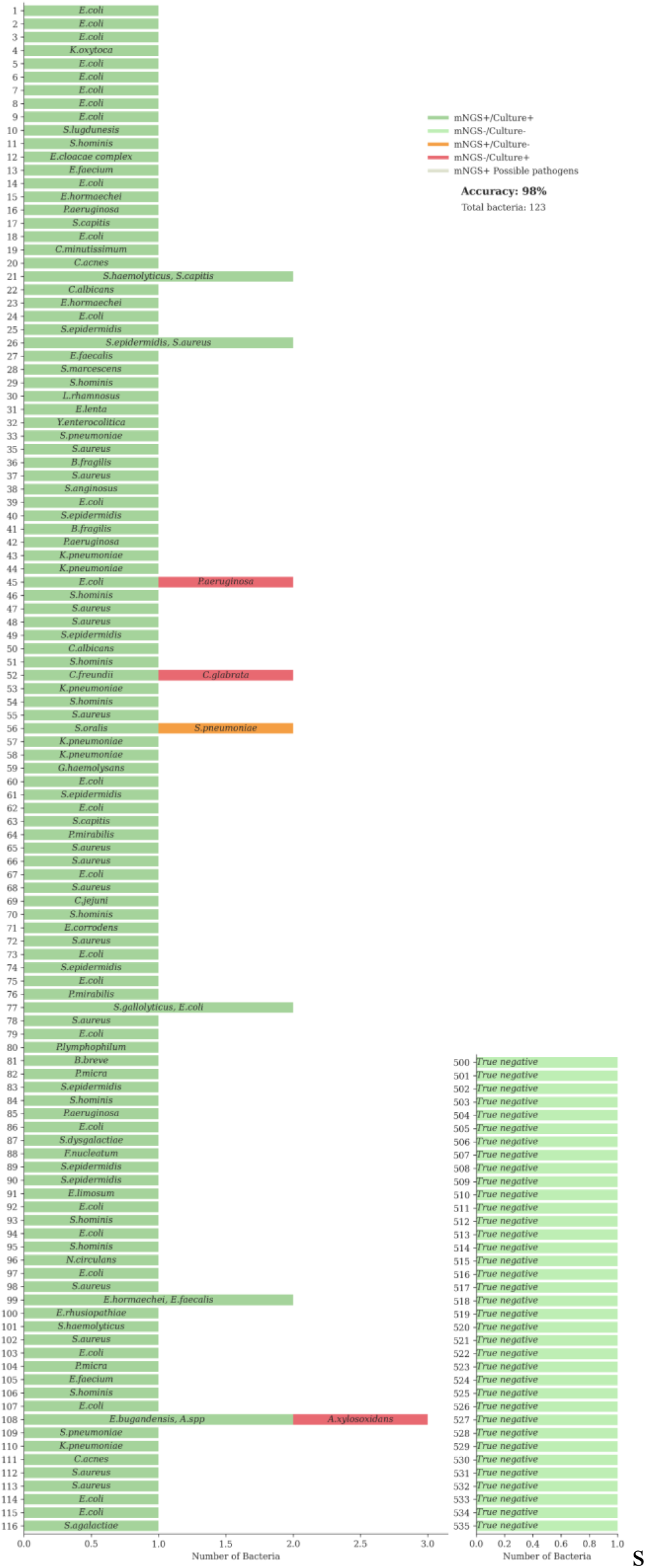
Overview of the pathogen identification. Benchmarking of the microbial species identification by mNGS compared to the routine clinical method (*definite MALDI*). Dark green bars represent species identified through both routine testing and mNGS (concordant). Light Green bars indicate the samples found to be negative through both routine testing and mNGS (concordant). Orange bars highlight those species uniquely identified by mNGS, and red bars denote species uniquely identified by routine testing. Grey indicates species identified by mNGS and confirmed with PCR. Species names are annotated in the bars, and the x-axis indicates the number.

The method also identified additional bacterial species that might be potential pathogens in five samples (1, 2, 5, 18, 39) that were missed by the *definite MALDI*. These additional species were *S. aureus* in 4 samples and *K. pneumoniae* in one sample (Table 1). The pathogen confirmed by the *definite MALDI* in all these samples was *E. coli*, which was also detected by mNGS. All these additional species were confirmed using qPCR with species-specific primers (Additional File 1: Table S7). The MLST analysis showed that the additional *S. aureus* detected by mNGS belonged to distinct sequence types and therefore may not be potential contaminants.

### mNGS showed 95% accuracy for AST prediction compared to the routine disk diffusion assay

Out of the 123 detected pathogens, phenotypic AST was performed on 120 isolates. A total of 1451 antibiotic-pathogen combinations were tested. The AST predictions from the mNGS, compared with the routine disk diffusion method, are summarized in Additional File 1: Fig S4. In AST prediction, the mNGS method achieved an accuracy of 95% (1382/1451), sensitivity of 88% (203/230), and specificity of 97% (1179/1221) (Fig 1c). The majority of false-negative results were observed with amoxicillin (n = 3), amoxicillin/clavulanic acid (n = 2), ampicillin (n = 2), trimethoprim/sulfamethoxazole (n = 4), and piperacillin/tazobactam (n = 5), mostly in *E. coli* (Fig 3). Similarly, most false-positive results occurred with amoxicillin/clavulanic acid (n = 9) and the cephalosporin class (n = 18) (Fig 3a).

**Fig 3:**
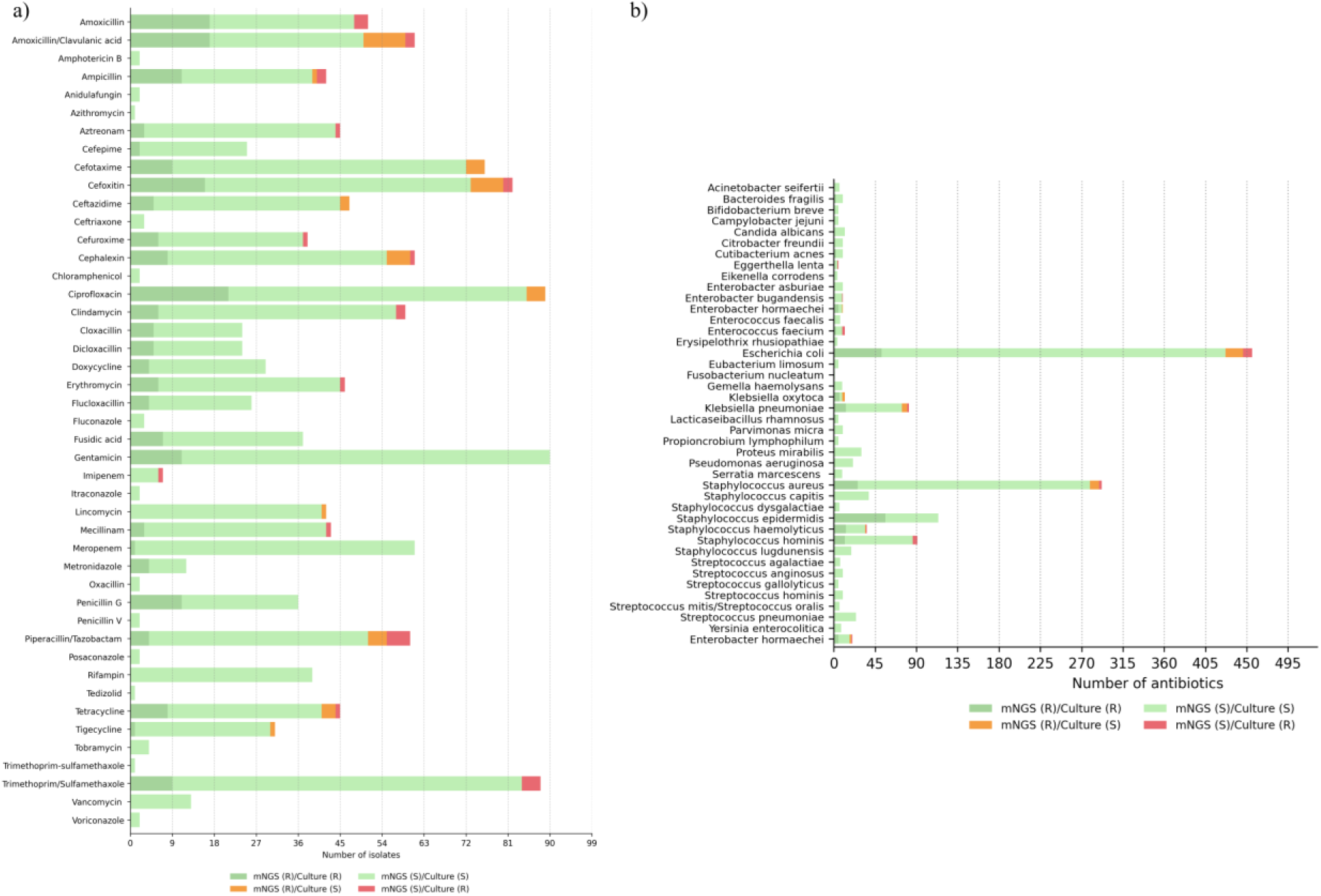
Overview of the antimicrobial susceptibility predictions. (a) Concordance between the mNGS results and the routine AST (disk diffusion) on the antibiotic level. (b) Concordance ordered by species. Dark green bars indicate that both routine AST and mNGS results suggest resistance, whereas light green bars indicate that both routine AST and mNGS indicate susceptibility. Red indicates cases where resistance was identified in routine AST, but no corresponding resistance mechanism was detected through mNGS (false negative). Orange bars denote detected ARGs without phenotypic resistance to the corresponding antibiotic (false positive).

Of the 28 *E. coli* samples, 13 were phenotypically susceptible to all tested antibiotics, and mNGS results confirmed this, as no relevant ARGs were detected in these samples. Two *E. coli* samples were resistant to only one antibiotic, whereas the others showed resistance to multiple antibiotics. *E. coli* sample 8 exhibited the highest resistance, being resistant to 11 of the 17 antibiotics tested, and mNGS accurately predicted the AST for 15 antibiotic combinations. In total, mNGS correctly predicted 427 antibiotic-pathogen combinations (52 TPs and 375 TNs) in *E. coli* samples (Fig 3b).

In *S. aureus*, five samples (out of 15) showed susceptibility to all tested antibiotics, which was confirmed by mNGS. mNGS achieved 265 correct predictions for *S. aureus*, including 26 TP and 239 TN, with only 13 errors (10 FPs and 3 FNs). 6 out of 10 *S. hominis* samples were phenotypically susceptible to all tested antibiotics, and the mNGS results showed the same. The remaining four samples were resistant to more than one antibiotic. mNGS results for *S. hominis* matched the phenotypic AST for 96 antibiotic-pathogen combinations, including 12 TPs and 84 TNs, with only 5 FNs. In *S. epidermidis,* only one sample was susceptible to all the tested antibiotics, as shown by both routine AST and mNGS, while the remaining samples were resistant to more than one antibiotic. mNGS correctly identified resistance or susceptibility for all 114 antibiotic combinations (56 TPs and 58 TNs), with no FPs or FNs detected. Similarly, in *K. pneumoniae*, three samples were phenotypically susceptible to all tested antibiotics, as confirmed by mNGS. In the remaining *K. pneumoniae* samples, mNGS made only eight incorrect predictions (six FPs and two FNs) (Fig 3b).

### Detection of diverse ARGs

β-Lactam resistance genes were the most frequently detected resistance determinants, including *ampC variants, TEM-1, SHV variants, ACT, OXY-1-4, SRT-2, cepA, and blaZ/PC1*. These ARGs were distributed across species, but most were detected in *E. coli*. Multidrug efflux pump-associated determinants were the second most abundant resistance category, frequently detected in *E. coli* and *S. epidermidis*. Detection of the methicillin resistance genes *mecA* and *mecR1* was primarily in *S. epidermidis*, with one sample each from *S. hominis* and *S. haemolyticus*. The majority of aminoglycoside resistance genes were also detected in *S. epidermidis*. Similarly, tetracycline resistance genes such as *tet38*, *tetK*, and *tetM* were detected in *S. epidermidis*, *S. hominis,* and *S. aureus* (Additional File 1: Fig S5).

### 94% of pathogens were identified in 30 minutes of sequencing

mNGS data collected at various time points after the start of the run showed that 94% (112) of detected species were identified within 30 minutes. Moreover, 79% (94) of species were identified after only 10 minutes of sequencing (Fig 4a). After 60 minutes of sequencing, 97% (115) of pathogens were detected, with only *C. albicans* (sample 50), *Campylobacter jejuni* (sample 69), and *E. faecium* (sample 105) being undetected. These were detected at 360, 600, and 1200 minutes, respectively (Additional File 1: Fig S6).

**Fig 4:**
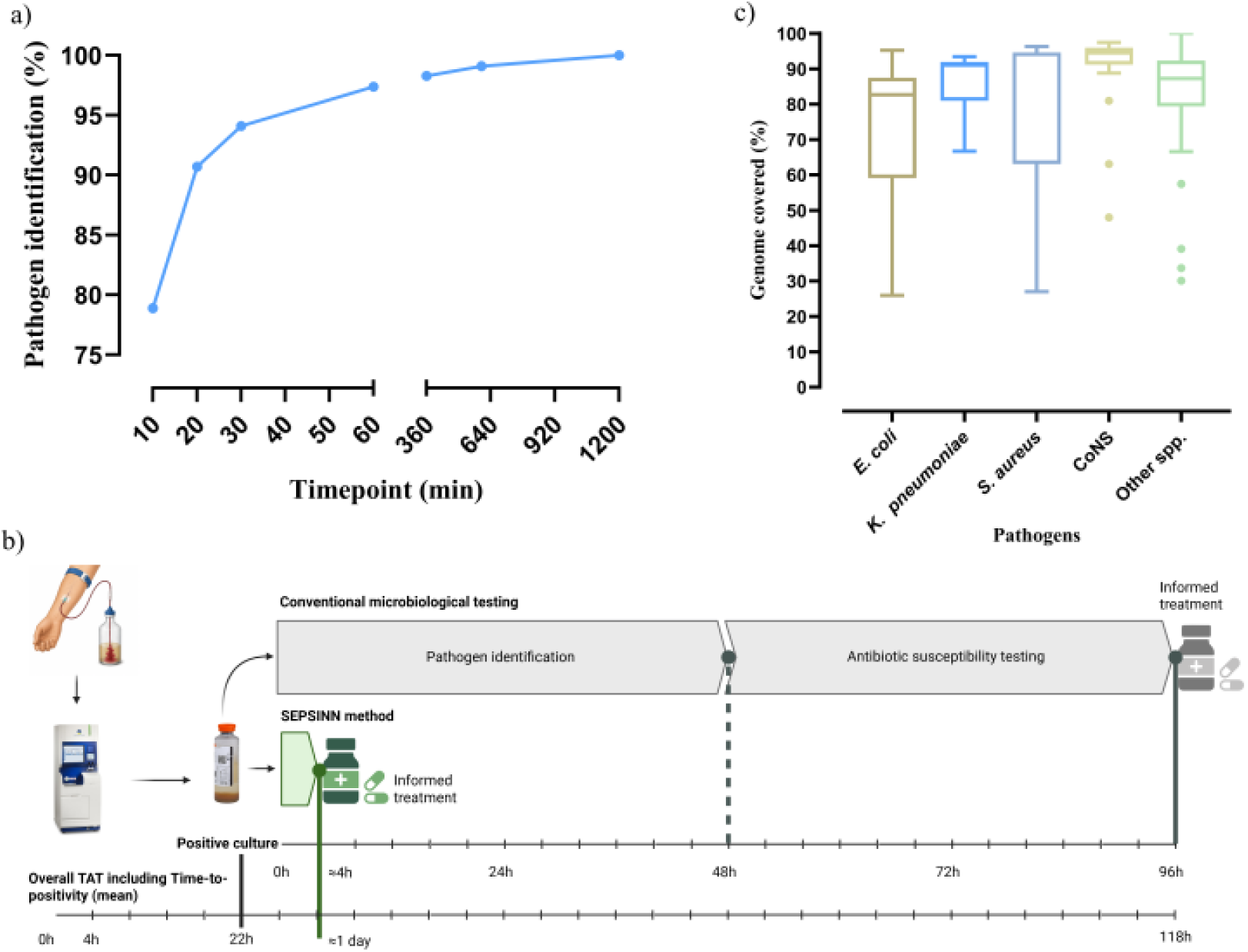
Analysis of the nanopore sequencing data showing. (a) Timeline for pathogen identification. Percentage of pathogen identified at different time points after the start of the sequencing run. (b) The timeline for pathogen identification and antibiotic resistance gene detection, using both conventional methods and mNGS. The total turnaround time using mNGS after the culture was flagged positive was approximately four hours. However, the mean time-to-positivity for blood cultures was 22 hours, and thus the total time to pathogen identification and antimicrobial susceptibility prediction from blood sample collection was approximately 1 day. (c) Genome coverage for the most frequently detected bacterial pathogens. Genome coverage across species is presented in groups, and CoNS included *S. epidermidis, S. haemolyticus, S. hominis, S. capitis,* and *S. lugdunensis.* Boxplots show the distribution of the genome coverage percentage. The boxes show the interquartile range (IQR); whiskers denote the range excluding outliers; points represent outliers. The line inside the box presents the median genome coverage.

### Approximately 4 hours of turnaround time from culture positivity to pathogen identification and ARG detection

The SEPSINN DNA extraction workflow, combined with nanopore sequencing, enabled the detection of both pathogens and ARGs from positive blood cultures within approximately 4 hours of sample receipt in the laboratory (Fig 4b). When the blood culture time-to-positivity is included (mean 22 hours), the overall turnaround time (TAT) is approximately one day. However, for samples with a quicker time-to-positivity, the TAT was < 12 hours. The TAT included blood culture incubation (≈ 22 hours), host DNA depletion (≈ 30 min), DNA extraction and purification (≈ 90 min), library preparation (≈ 60 min), nanopore sequencing, and real-time bioinformatic analysis (≈ 60 min).

### 90% of the median genome coverage was observed using mNGS

Genome coverage varied between different species groups. The median genome coverage for all the species was approximately 90%. *E. coli* exhibited a median genome coverage of 83%, with only four samples showing a genome coverage of less than 50%. *K. pneumoniae* consistently showed higher genome coverage, with a median of 91%, and only one sample had a comparatively lower genome coverage of 67%. In *S. aureus*, a median genome coverage of over 94% was observed, with only three samples showing coverage below 90%. Similarly, CoNS species had a median genome coverage of over 94%, with the majority of samples showing coverage above 90%. The genome coverage for all other species was grouped and observed to have a median of 87% (Fig 4c).

## Discussion

In this study, we have shown that our in-house-developed host depletion and DNA extraction method (SEPSINN), when combined with real-time nanopore sequencing and data analysis, achieved high accuracy in identifying pathogens and profiling their resistance within approximately four hours of positive blood cultures. Moreover, using SEPSINN, the pathogen and ARGs were detected approximately one day after receipt of the blood sample in the laboratory. The method performed on par with clinical routine diagnosis, which used MALDI (purified colonies) for pathogen ID and disk diffusion for AST testing. However, the SEPSINN mNGS method outperformed the *initial MALDI*, which was performed on positive blood cultures without isolating pure bacterial colonies. Furthermore, we used an unselective sampling process, ensuring a continuous supply of samples from clinical routine without selection bias.

Blood culture is considered the gold standard for diagnosing BSIs [36,37]. However, isolating bacterial DNA from blood cultures is challenging because human cells outnumber microbial cells, have a larger genome size, and contain about 1500 times more DNA per cell [38–40]. To enhance bacterial growth and neutralize the effects of antibiotics, blood culture bottles are supplemented with various additives, such as sodium polyanetholsulfonate (SPS), gelatin, resins, and charcoal[18]. These additives, specifically SPS, which is usually co-purified with the extracted DNA, are strong inhibitors of downstream molecular-based processes such as PCR and mNGS [19,41]. Most previous metagenomic studies have used commercially available kits for DNA extraction from blood culture [12,14,16,42–47]. Specifically, the BiOstic bacteremia kit is most commonly used, including in our previous studies involving spiked blood cultures [12,13,24,48]. While the BiOstic kit performed well, it lacks host DNA depletion, which is important for increasing microbial sequencing data and streamlining analysis. Host DNA depletion will be especially beneficial when the microbial load is low, particularly in polymicrobial samples where one species is less abundant than the others. Depleting host DNA is also essential for obtaining sufficient microbial mNGS data for precise AMR predictions [49]. The SEPSINN method successfully isolated bacterial DNA from blood cultures while efficiently depleting host DNA and removing inhibitors. Up to 1000-fold depletion of host DNA was observed relative to non-depleted samples. Previous studies, including ours, have reported greater fold depletion with saponin- and nuclease-based methods; however, those studies used spiked samples, whereas this study used blood cultures from patients with BSI [20,23,24].

The SEPSINN mNGS method detected pathogens at the species level. All monomicrobial infections, including fungal infections, were identified to the species level. Among the seven polymicrobial samples, all pathogens were detected in four, and one pathogen was detected in the remaining three. Previous studies have shown that nanopore sequencing can detect monomicrobial samples at the genus level, whereas polymicrobial samples are challenging to identify [16,43,45,47,50]. However, most of these studies focused only on bacterial species identification; in the current study, all pathogens, including fungi, were detected. In addition, the mNGS method identified five additional pathogens compared with the *definite MADI*. The identification of these additional pathogens was confirmed with qPCR using species-specific primers. A recent study reported similar results, showing that a sequencing-based method can detect additional fastidious and unculturable microbes [12,23].

An important aspect of using an mNGS method in clinical diagnostics is its ability to predict antimicrobial susceptibility. For AST predictions, our method achieved 95% accuracy, 97% specificity, and 88% sensitivity across 1451 antibiotic combinations. These results showed greater concordance with the phenotypic AST than those reported in previously published studies. Harris et al.[16] achieved 87% accuracy, Azami et al.[43] 76%, and Gu et al.[42] 80%. A recent study by Govender et al. reported 88% sensitivity and 93% specificity for AST predictions [12]. Most of these previous studies have used the R9.4 flow cells from ONT, which rely on older chemistry and outdated basecalling and demultiplexing algorithms. However, the current study used the latest R10 chemistry, which offers a lower error rate, higher Q score and accuracy, and updated machine-learning-based algorithms for basecalling [51,52]. Additionally, the current study predicted AST using a diverse sample set comprising aerobic and anaerobic bacteria. However, most previous studies have focused on specific species for resistance prediction [10,11], while others also perform whole-genome amplification before sequencing, which can introduce bias in AMR prediction [16,53]. Most false-negative results were observed with amoxicillin, amoxicillin/clavulanic acid, ampicillin, trimethoprim/sulfamethoxazole, and piperacillin/tazobactam. False-negative results for these antibiotics in *E. coli* have also been observed previously [23,47]. Resistance to these antibiotics usually results from multiple genetic elements, including chromosomal mutations, overexpression of efflux pumps, and plasmid-mediated resistance [54]. It is difficult to detect these mutations or overexpressed efflux pumps and correlate them with the phenotypic AST, making the prediction challenging. Similarly, most false-positive predictions were observed with cephalosporins for *E. coli* and *K. pneumoniae*, and with amoxicillin/clavulanic acid for *S. aureus*. The percentage of the bacterial genome sequenced for each species is also important for accurate AST prediction [55]. Our method generally achieved good genome coverage across most samples; however, some *E. coli* samples had coverage below 50, which could explain why some resistance predictions were missing.

The current clinical routine diagnostic approach to BSI takes 2–4 days, necessitating a lengthy regimen of empirical antibiotic treatment in severe cases. mNGS-based methods have the potential to reduce the TAT from days to hours. However, workflows based on Illumina sequencing remain time-consuming, with a turnaround time of 23 to 59 hours, comparable to that of current culture-based methods [44]. Nanopore sequencing can overcome these limitations, as it is fast and provides real-time data. Previously published studies have reported TATs of 6–16 hours for pathogen identification and AMR prediction in BSIs using nanopore sequencing [16,43]. Others have reported comparatively shorter TATs of 4.4 hours [47] and 6 hours [12]. However, these studies either focus on selective resistance genes or pathogens for AST predictions. In our study, using an unbiased approach to select all samples, we observed an approximate 4-hour TAT for species identification and AMR predictions.

The results show a mean TTP of about 22 hours; however, there is a significant difference in TTP between aerobic (≈17 hours) and anaerobic (≈66 hours) bacteria. But there is no significant difference in turnaround times for downstream DNA extraction, sequencing, and bioinformatics analysis. This means that for aerobic blood cultures, information for informed decision-making can be obtained in about 21 hours. Because blood culture is performed separately for aerobic and anaerobic bacteria in two culture bottles, a stratified metagenomic approach is possible. This is especially significant, as the vast majority of BSI pathogens are aerobic, and only a small percentage (ca. 5%) are anaerobic [56–58]. It would therefore be beneficial to analyze the first positive bottle and then perform less laborious procedures to confirm that subsequent bottles contain the same or additional pathogens.

While the method was tested on a diverse sample set comprising 109 aerobic and 11 anaerobic bacteria, along with three fungi, the samples originated from a single geographical region with relatively low AMR prevalence, which may limit generalizability to high-resistance settings. The method showed improved AMR prediction accuracy compared with previous studies; however, there remains a need to understand the diverse resistance mechanisms underlying some antibiotic-drug combinations [42]. The current method is manual and requires bioinformatics expertise to analyze the generated data, hindering its implementation in clinical settings. The method was evaluated using positive blood cultures; however, to further reduce time and eliminate the longer time-to-positivity of blood culture, efforts should focus on sequencing whole blood samples directly from the patient [48].

### Conclusions

The results demonstrate the effectiveness of our in-house developed host depletion and DNA extraction method (SEPSINN), enhanced by nanopore sequencing and real-time data analysis, for the rapid diagnosis of BSIs in clinical settings. Overall, the method achieved 98% pathogen identification accuracy and 95% antibiotic susceptibility prediction accuracy, with host depletion up to 1000-fold. This approach achieves an overall accuracy exceeding 90%, which is typically essential for implementing new diagnostic techniques in clinical microbiology [59,60]. Furthermore, these mNGS techniques can provide deeper insights into the mechanisms of resistance. The method took approximately 4 hours for pathogen detection and antimicrobial resistance profiling after culture positivity. Since sepsis is time-critical, achieving a turnaround time of approximately 24 hours from sample receipt in the laboratory, and potentially even faster for aerobic blood cultures, could prevent millions of unnecessary antibiotic doses of empirical broad-spectrum antibiotics each year before initiating personalized, targeted therapy against an identified pathogen. It will also promote better clinical decisions in managing BSI/sepsis and support antibiotic and diagnostic stewardship.

## Declarations

### Ethics approval and consent to participate

The study was approved by the regional medical and health research ethics committee (REC Helse Sør-Øst) (691177). The study was conducted in accordance with the local legislation and institutional requirements. The ethics committee waived the requirement for written informed consent for participation by participants or their legal guardians/next of kin, as the samples were anonymized and no patient-related information was collected.

### Consent for publication

“Not applicable”

### Availability of data and materials

The dataset supporting the conclusions of this article is available in the NCBI Sequence Read Archive repository, with BioProject ID PRJNA1483694, https://www.ncbi.nlm.nih.gov

### Declaration of competing interests

The authors declare no competing interests that could have influenced the work reported in this paper.

### Funding

The study was financially supported by the OH-AMR-Diag project, funded by the Research Council of Norway (Project number 336420).

### Author contributions

Jawad Ali: Conceptualization, Formal analysis, Investigation, Methodology, Visualization, Writing—original draft, Writing—review and editing. Anurag Basavaraj Bellankimath: Methodology, Formal analysis, Visualization, Investigation, Writing—review and editing. Silje Therese Opgård: Methodology, Formal analysis, Investigation, Writing—review and editing. Emil Varman Manivannan: Methodology, Investigation, Writing—review and editing. Gunnar Skov Simonsen: Conceptualization, Project administration, Writing—review and editing. Rafi Ahmad: Conceptualization, Formal analysis, Project administration, Funding acquisition, Supervision, Writing—original draft, Writing—review and editing. All authors contributed to the manuscript and approved the submission.

## Notes

### Competing Interest Statement

The authors have declared no competing interest.

