## Additional File 1 for "From culture to clarity in four hours: accelerating clinical management of bloodstream infections using metagenomics"

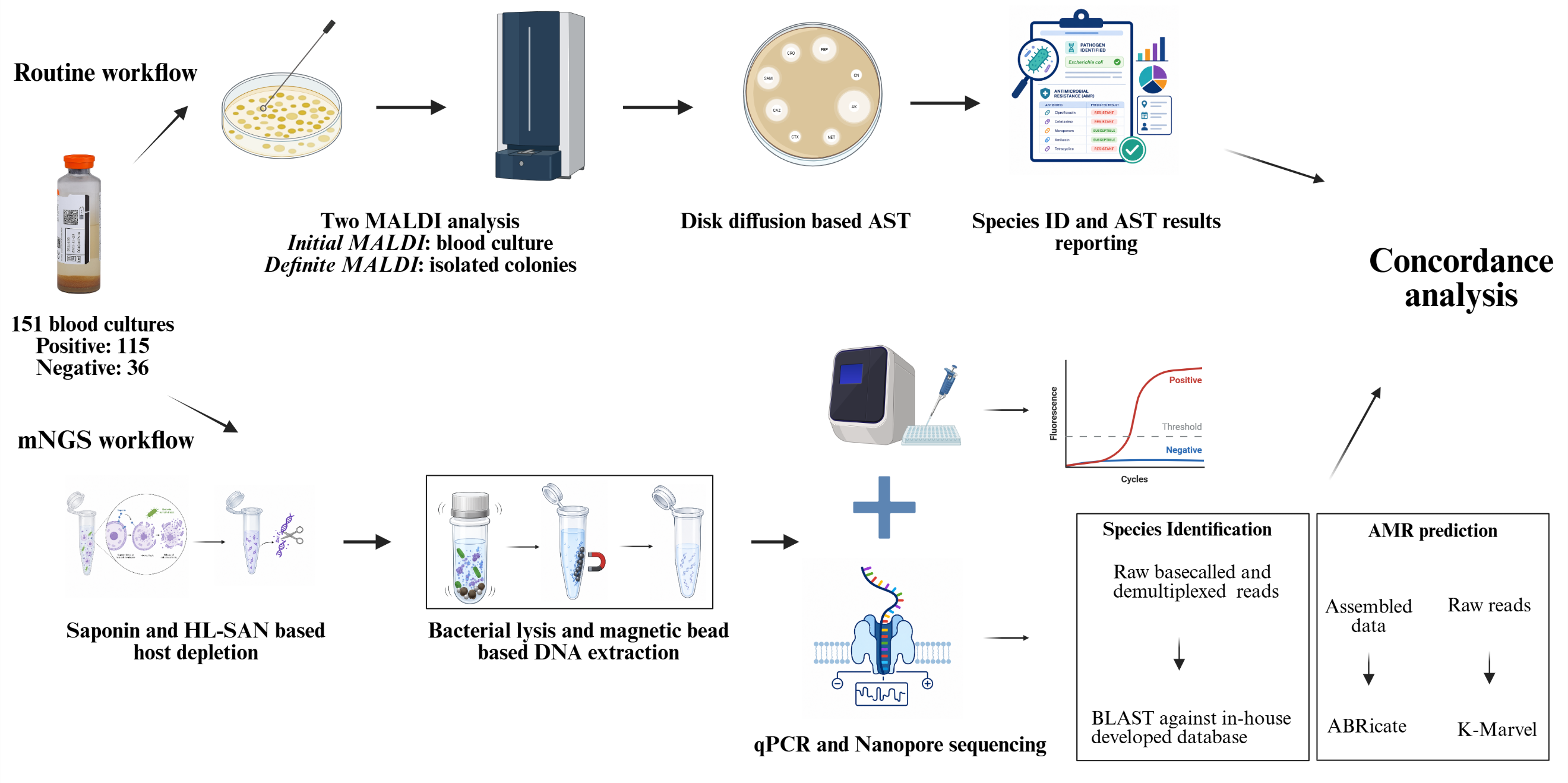

**Fig S1: Overview of experimental design.** Blood cultures were processed through both the routine diagnosis and mNGS workflows. In routine workflow, a positive blood culture is collected, an *initial MALDI* is performed on the culture, followed by subculturing and performing a *definite MALDI* on the isolated colonies. For the mNGS workflow, 1–2 mL of blood cultures were processed for host depletion using saponin and HL-SAN nuclease, followed by DNA extraction using the NAXTRA bead-based method. After DNA extraction, nanopore sequencing, and qPCR, data analysis was performed. The mNGS-based species and antimicrobial susceptibility predictions were analyzed for concordance with the routine-based results.

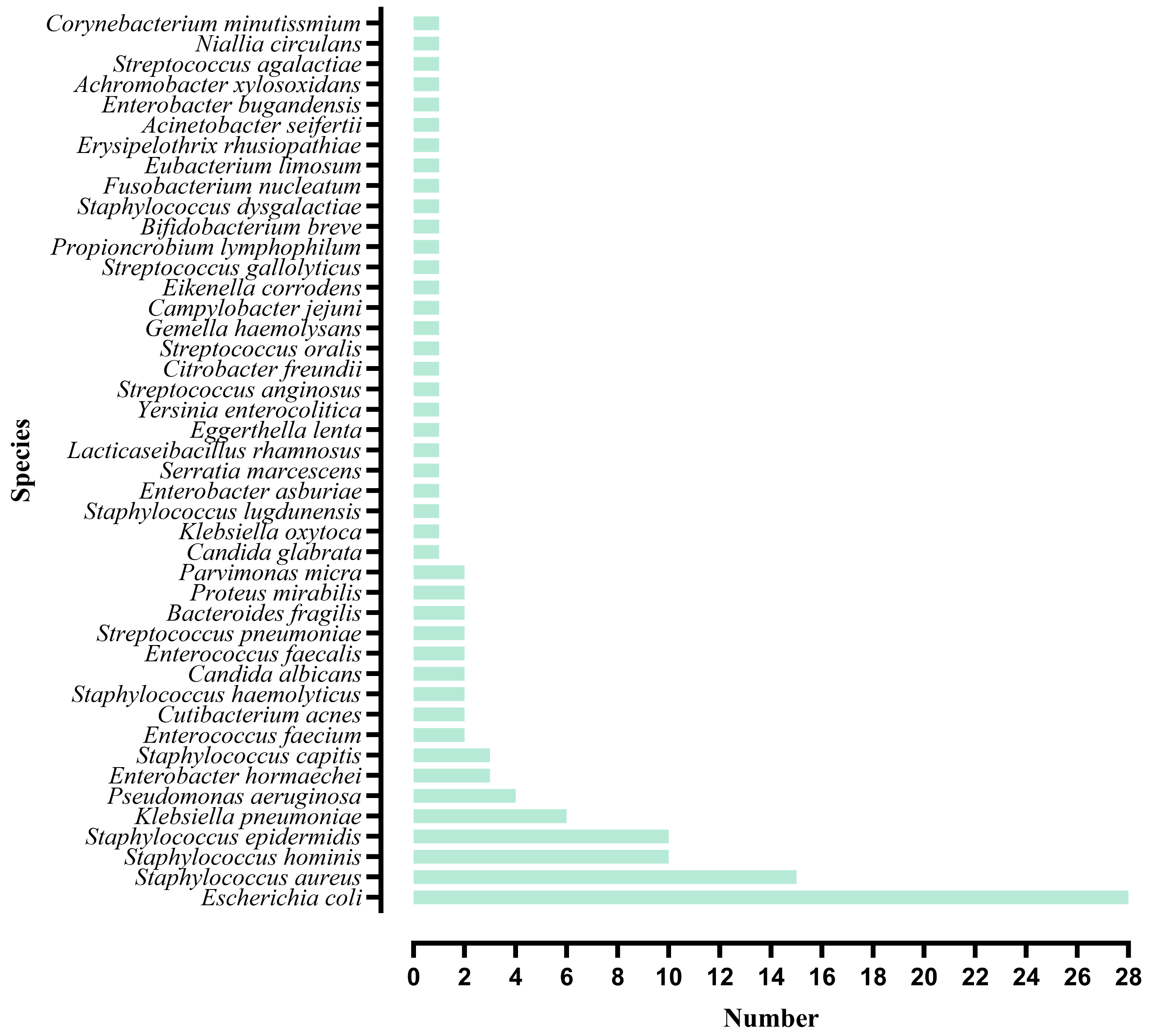

b)

a)

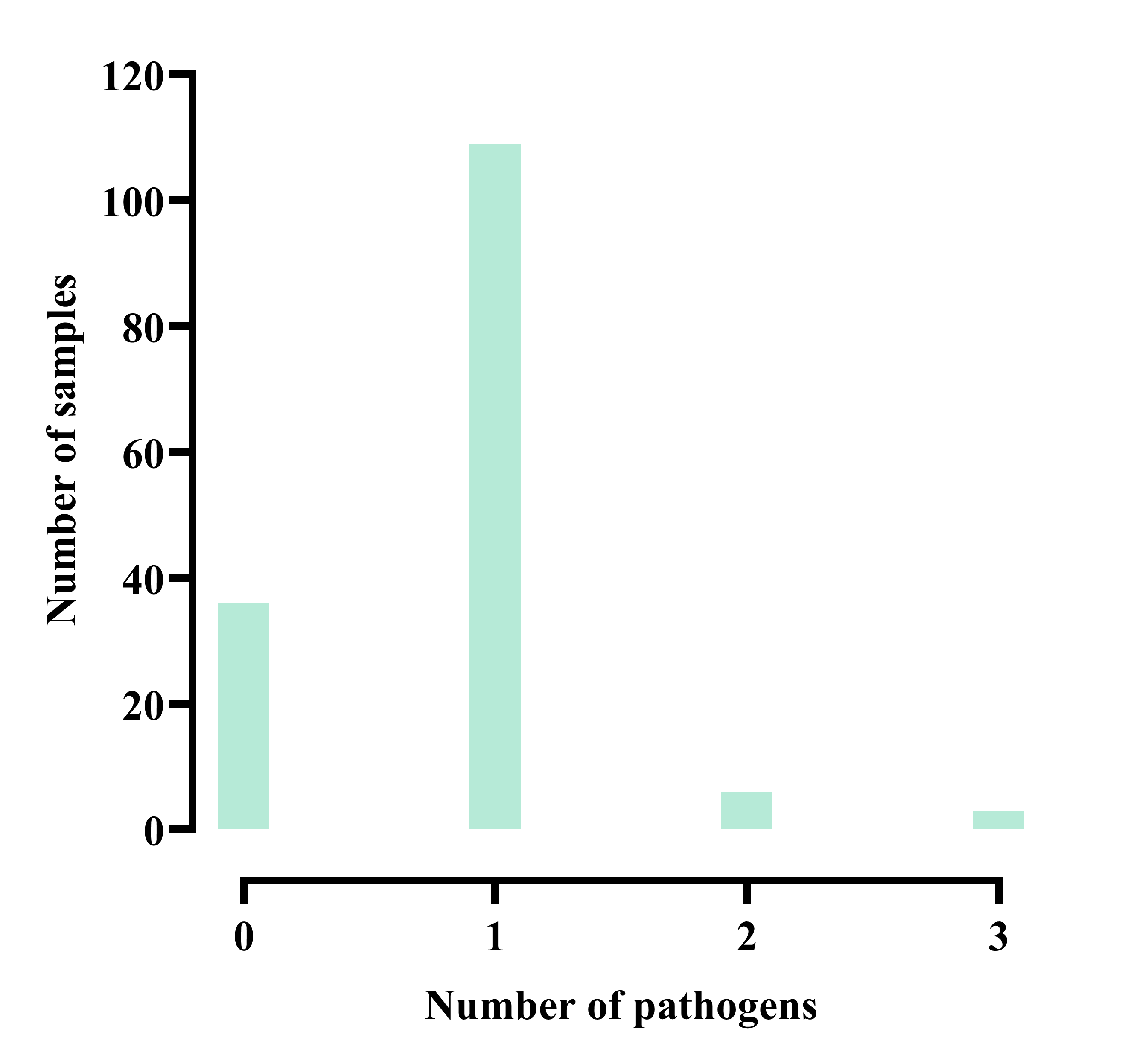

**Fig S2:** **Summary of bacterial pathogens identified in blood cultures through routine clinical MALDI detection.** (a) Diverse microbial species were observed across the samples, including aerobic and anaerobic bacteria and fungi (b) Histogram showing the number of pathogens in different samples that were detected by definite MALDI.

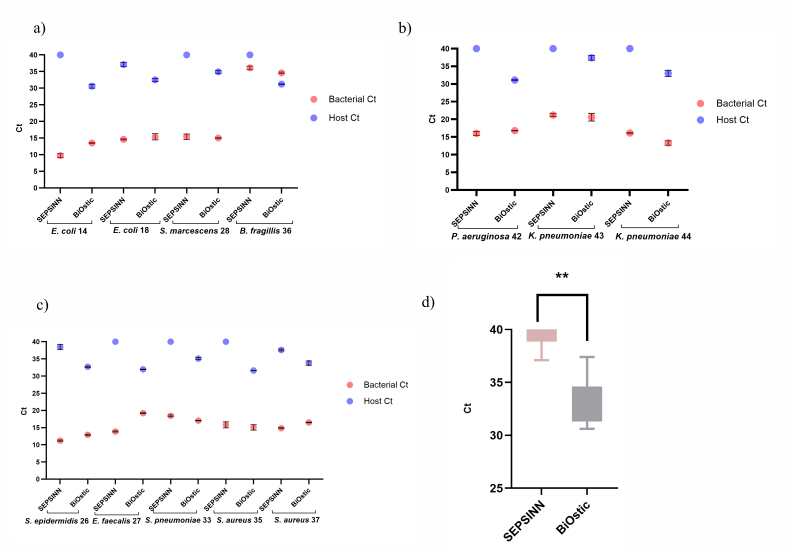

**Fig S3: qPCR Ct values showing the comparison of the host DNA depletion and bacterial DNA recovery using SEPSINN and BiOstic extraction methods.** (a) and (b) Gram-negative pathogens (c) Gram-positive pathogens (d) Statistical significance of the host DNA depletion between SEPSINN and BiOstic. ** = statistical significance

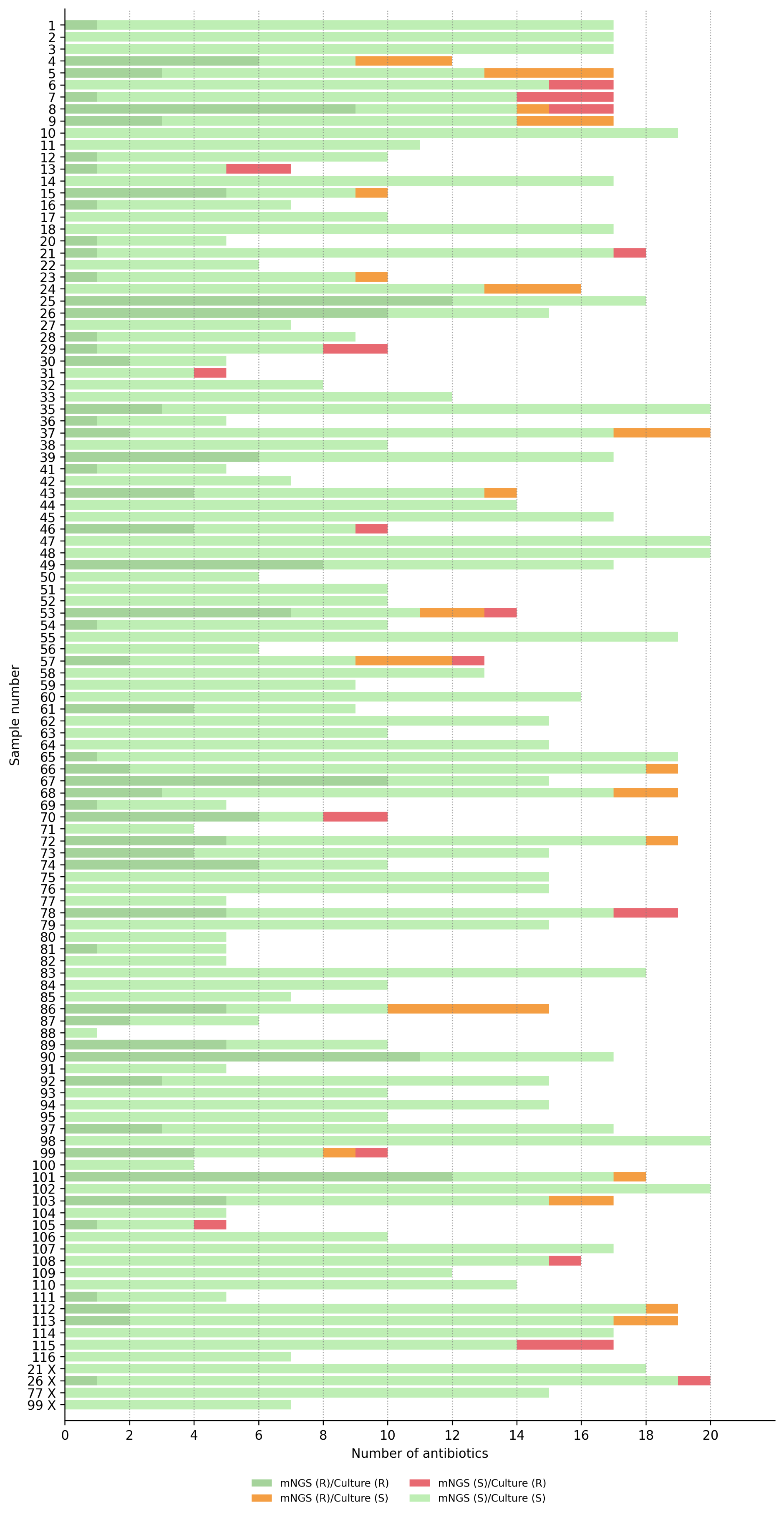

**Fig S4: Overview of antimicrobial susceptibility benchmarking.** The results for AST predictions from mNGS compared with the reference method (disk diffusion) are shown for each sample. The dark green bar indicates resistance by both mNGS and the routine method, whereas the light green bar shows susceptibility by both. Orange bars indicate the detection of resistance genes by mNGS, but no phenotypic resistance was observed by the routine method. Red bar demonstrates phenotypic resistance by the routine method, but no corresponding resistance determinants were detected by mNGS. “X” demonstrates polymicrobial samples.

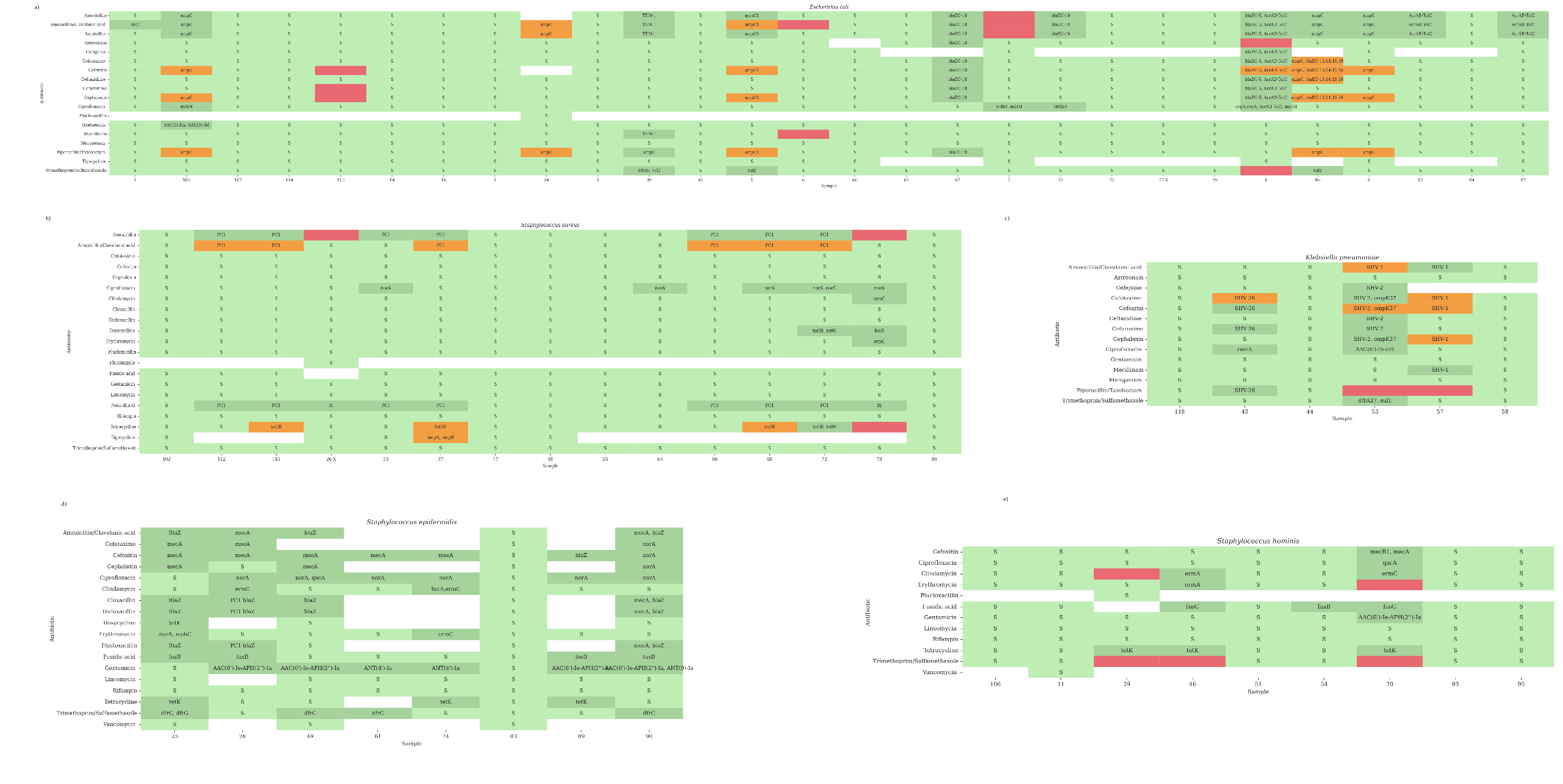

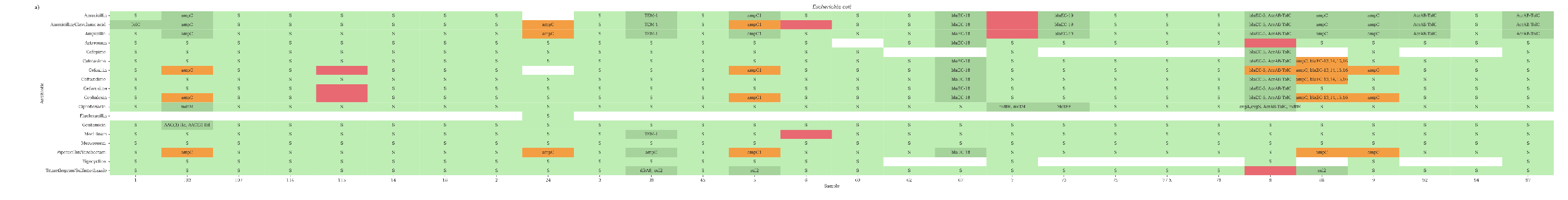

**Fig S5:** **Detailed overview of the AST benchmarking.** Heatmaps illustrating the concordance between routine antimicrobial susceptibility testing and antimicrobial resistance predictions derived from metagenomic next-generation sequencing (mNGS) and detected antimicrobial resistance genes (ARGs). Results are grouped by pathogen, with antibiotic-specific susceptibility and resistance profiles shown for each sample. Green cells indicate concordant results between phenotypic AST and mNGS predictions: light green denotes susceptibility by both methods, whereas dark green denotes resistance identified by both methods. Cells are annotated with **S** (susceptible), or the corresponding ARG when a resistance determinant was detected. Red cells indicate phenotypic resistance detected by AST, but no corresponding resistance mechanism is identified by mNGS (false-negative genotypic prediction). Orange cells indicate ARG detection in the absence of phenotypic resistance (genotype–phenotype discordance) and are annotated with the detected ARG(s). Antimicrobial resistance predictions were evaluated only for antibiotics represented in both the phenotypic susceptibility dataset and the ARG reference database. (a) *E. coli* (b) *S. aureus* (c) *K. pneumoniae* (d) *S. epidermidis* (e) *S. hominis*.

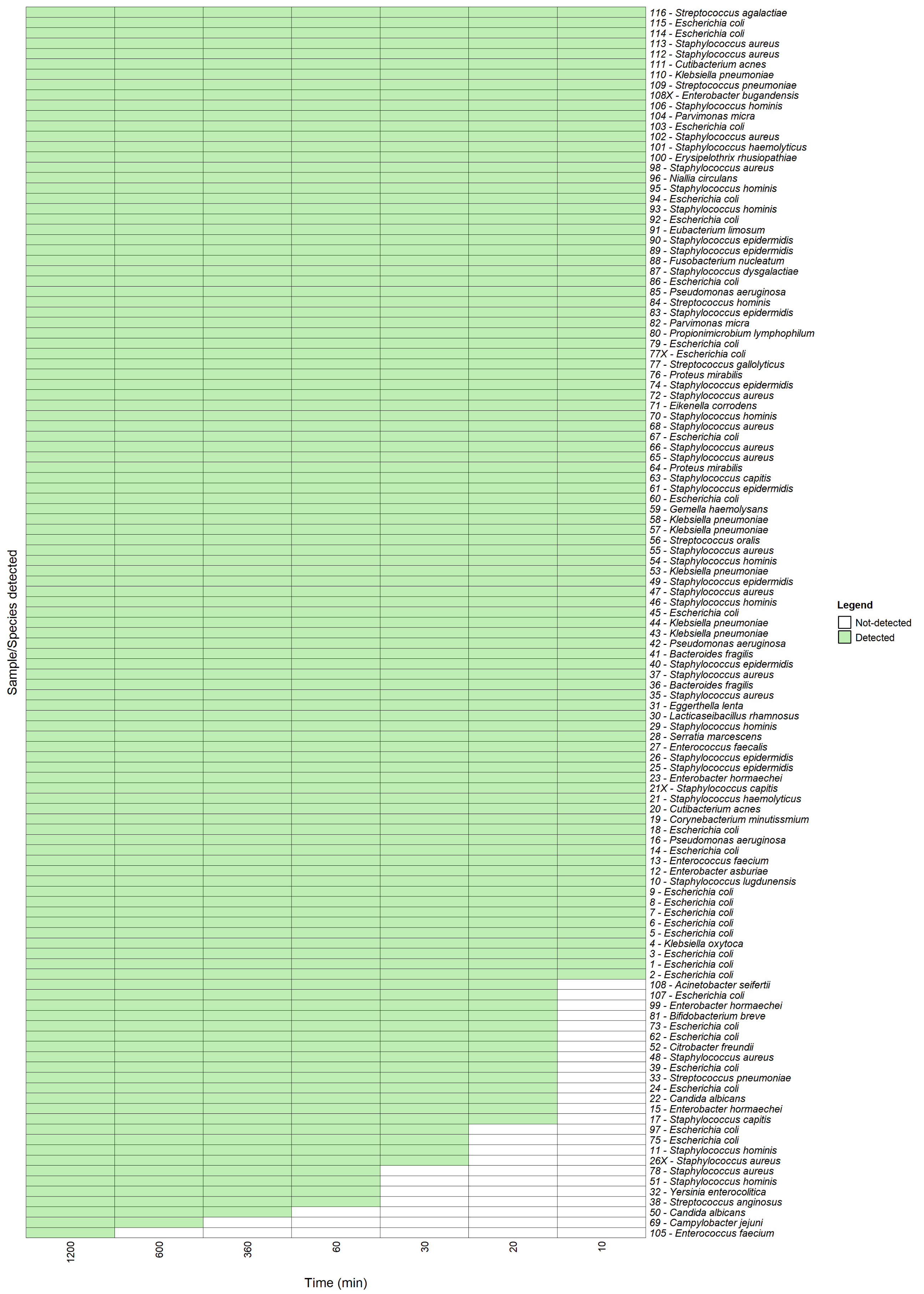

**Fig S6: Timeline for pathogen identification using mNGS data.** Identification of the bacterial pathogens at different time points after the start of the sequencing run. Each row represents an individual microbial species, along with the sample numbers, and each column represents the elapsed time after the start of the sequencing run.

**Table S1:** List of species-specific primers used for qPCR-based detection of bacterial pathogens and host DNA. FP= Forward primer, RP= Reverse primer

| **Target species** | **Target gene** | **Amplicon size (bp)** | **Primers sequences** | **References** |
| --- | --- | --- | --- | --- |
| Human | β-Actin | ~100 | FP: CGGCCTTGGAGTGTGTATTAAGTA RP: TGCAAAGAACACGGCTAAGTGT | [1] |
| *E. coli* | *UspA* | ~850 | FP: CCGATACGCTGCCAATCAGT  RP: ACGCAGACCGTAGGCCAGAT | [2] |
| *P. aeruginosa* | *phzA2* | ~325 | FP: GTTTACCGACAACCTGGAA  RP: GCAATAGCCCTGCGGATAC | [3] |
| *S. aureus* | *Nuc* | ~65 | FP: GGGTTGATACGCCAGAAACG  RP: TGATGCTTCTTTGCCAAATGG | [4] |
| *K. pneumoniae* | *Khe* | ~486 | FP: TGATTGCATTCGCCACTGG  RP: GGTCAACCCAACGATCCTG | [5] |
| *S. epidermidis* | *altE* | ~682 | FP: CAACTGCTCAACCGAGAACA  RP: TTTGTAGATGTTGTGCCCCA | [6] |
| *E. faecalis* | *GroES* | ~185 | FP: GGAATTGTTCTTGCATCCGT  RP: ACAATTAAGTATTCTACGCC | [7] |
| *S. marcescens* | *luxS* | ~516 | FP: TGCCTGGAAAGCGGCGATGG  RP: CGCCAGCTCGTCGTTGTGGT | [8] |
| *S. pneumoniae* | *Spn9802* | ~157 | FP: AGTCGTTCCAAGGTAACAAGTCT  RP: ACCAACTCGACCACCTCTTT | [9] |
| *B. fragilis* | *RecA* |  | FP: CAGTCAAGGCGGCTACAGAG  RP: CAGTTTAGCCGCATAGAAGC | [10] |

**Table S3:** qPCR results for human and bacterial DNA from blood cultures with and without host DNA depletion. Host depletion and bacterial DNA recovery in the depleted samples were calculated using ΔCt values normalized to the non-depleted controls. Host DNA amplification in most depleted samples was above the detection threshold (40 PCR cycles). However, to calculate the host depletion efficiency, we have used 40 as the Ct value for those samples.

| **Sample number** | **MALDI ID** | **SEPSINN ID** | **Sample treatment** | **Human qPCR assay (Avg. Ct)** | **Human DNA depletion (ΔCt)** | **Bacterial species-specific qPCR (Avg. Ct)** | **Bacterial gain/loss (ΔCt)** |
| --- | --- | --- | --- | --- | --- | --- | --- |
| 2 | *E. coli* | *E. coli* | Depleted | > 40.0 | 7.31 (~10^2^) | 15.0 | 1.73 |
|  |  |  | Non-depleted | 32.7 |  | 16.8 |  |
| 3 | *E. coli* | *E. coli* | Depleted | > 40.0 | 5.82 (~10) | 14.7 | 3.30 |
|  |  |  | Non-depleted | 34.2 |  | 18.0 |  |
| 5 | *E. coli* | *E. coli* | Depleted | > 40.0 | 6 (~10^2^) | 16.9 | 2.1 |
|  |  |  | Non-depleted | 34.0 |  | 14.8 |  |
| 6 | *E. coli* | *E. coli* | Depleted | > 40.0 | 7.16 (~10^2^) | 13.9 | 3.47 |
|  |  |  | Non-depleted | 32.8 |  | 17.4 |  |
| 7 | *E. coli* | *E. coli* | Depleted | > 40.0 | 6.76 (~10^2^) | 15.1 | 2.36 |
|  |  |  | Non-depleted | 33.2 |  | 17.5 |  |
| 8 | *E. coli* | *E. coli* | Depleted | > 40.0 | 6.69 (~10^2^) | 15.4 | 3.40 |
|  |  |  | Non-depleted | 33.3 |  | 18.8 |  |
| 9 | *E. coli* | *E. coli* | Depleted | > 40.0 | 0.60 (~2) | 15.4 | 1.60 |
|  |  |  | Non-depleted | 39.4 |  | 13.8 |  |
| 14 | *E. coli* | *E. coli* | Depleted | > 40.0 | 9.7 (~10^3^) | 13.7 | 0.10 |
|  |  |  | Non-depleted | 30.3 |  | 13.8 |  |
| 16 | *P. aeruginosa* | *P. aeruginosa* | Depleted | 39.0 | 6.5 (~10^2^) | 19.3 | 0.70 |
|  |  |  | Non-depleted | 32.5 |  | 20.0 |  |
| 18 | *E. coli* | *E. coli* | Depleted | 37.6 | 6.36 (~10^2^) | 14.8 | 0.49 |
|  |  |  | Non-depleted | 31.2 |  | 14.3 |  |

**Table S4:** DNA yield and qPCR Ct values for host and bacterial primers for samples using both the SEPSINN and BiOstic extraction methods. STD= Standard deviation

| **Sample number** | **Species** | **Extraction method** | **DNA yield (ng)** | **Bacterial Ct mean + STD** | **Host Ct mean**  **+ STD** |
| --- | --- | --- | --- | --- | --- |
| 14 | *E. coli* | SEPSINN | 34 400 | 9.7 ± 0.6 | 40.0 |
|  |  | BiOstic | 28 900 | 13.5 ± 0.2 | 30.6 ± 0.6 |
| 18 | *E. coli* | SEPSINN | 33 800 | 14.6 ± 0.2 | 37.1 ± 0.6 |
|  |  | BiOstic | 20 700 | 15.4 ± 0.9 | 32.5 ± 0.5 |
| 26 | *S. epidermidis* | SEPSINN | 9 450 | 11.2 ± 0.2 | 38.5 ± 0.7 |
|  |  | BiOstic | 43 200 | 12.9 ± 0.2 | 32.7 ± 0.2 |
| 27 | *E. faecalis* | SEPSINN | 88 00 | 13.9 ± 0.2 | 40.0 |
|  |  | BiOstic | 32 700 | 19.2 ±0.1 | 32.0 ± 0.1 |
| 28 | *S. marcescens* | SEPSINN | 45 000 | 15.4 ± 0.8 | 40.0 |
|  |  | BiOstic | 36 900 | 15.0 ± 0.2 | 34.9 ± 0.4 |
| 33 | *S. pneumoniae* | SEPSINN | 10 600 | 18.4 ± 0.3 | 40.0 |
|  |  | BiOstic | 13 200 | 17.0 ± 0.1 | 35.1 ± 0.3 |
| 35 | *S. aureus* | SEPSINN | 4 640 | 15.8 ± 0.9 | 40.0 |
|  |  | BiOstic | 13 700 | 15.1 ± 0.8 | 31.6 ± 0.0 |
| 36 | *B. fragilis* | SEPSINN | 63 000 | 36.1 ± 0.6 | 40.0 |
|  |  | BiOstic | 50 500 | 34.6 ± 0.3 | 31.2 ± 0.1 |
| 37 | *S. aureus* | SEPSINN | 5 280 | 14.8 ± 0.2 | 37.6 ± 0.3 |
|  |  | BiOstic | 27 000 | 16.5 ± 0.1 | 33.8 ± 0.6 |
| 42 | *P. aeruginosa* | SEPSINN | 22 800 | 16.0 ± 0.6 | 40.0 |
|  |  | BiOstic | 27 800 | 16.8 ±0.1 | 31.3 ± 0.2 |
| 43 | *K. pneumoniae* | SEPSINN | 19 200 | 21.2 ± 0.4 | 40.0 |
|  |  | BiOstic | 18 700 | 20.6 ± 1.1 | 37.4 ± 0.7 |
| 44 | *K. pneumoniae* | SEPSINN | 29 700 | 16.1 ± 0.2 | 40.0 |
|  |  | BiOstic | 31 600 | 13.3 ± 0.6 | 33.0 ± 0.9 |

|  | Prevalence threshold | Sensitivity | Specificity | Positive predictive value (PPV) | Negative predictive Value (NPV) | Accuracy |
| --- | --- | --- | --- | --- | --- | --- |
| Pathogen ID | 98% (156/159) CI 96-100% | 98% (120/123)  CI 95-100% | 97%  (36/37)  CI 92-100% | 99%  (120/121) CI 98-100% | 92%  (36/39)  CI 84-100% | 98% (156/160)  CI 95-100% |
| AST |  | 88% (203/230)  CI 84-92% | 97% (1179/1221) CI 96-98% | 83% (203/245) CI 78-88% | 98% (1179/1206) CI 97-99% | 95% (1382/1451) CI 94-96% |

**Table S6: An overview of the scoring matrices obtained by comparing mNGS (SEPSINN) results with routine MALDI/disk diffusion-based testing results.** Metrics for agreement between metagenomics results and routine pathogen identification and antibiotic susceptibility testing (AST) are shown on a pathogen/antibiotic basis. Each metric is expressed as a percentage and accompanied by the absolute numbers in parentheses. The 95% confidence interval (CI) is provided for each metric as a percentage.

| **Sample ID** | **Target species** | **Ct values** |
| --- | --- | --- |
| 1 | *S. aureus* | 25 |
| 2 | *S. aureus* | 27 |
| 5 | *K. pneumoniae* | 32 |
| 18 | *S. aureus* | 34 |
| 39 | *S. aureus* | 29 |

**Table S7:** qPCR Ct values for the confirmation of additional species detected through mNGS.
